# Understanding the transcriptional response to ER stress in Chinese hamster ovary cells using multiplexed single cell RNA-seq

**DOI:** 10.1101/2022.03.31.486542

**Authors:** Ioanna Tzani, Marina Castro-Rivadeneyra, Stefano Boi, Colin Clarke

## Abstract

Single cell RNA-seq (scRNA-seq) has recently been shown to provide a powerful method for the analysis of transcriptional heterogeneity in Chinese hamster ovary (CHO) cells. A potential drawback of current scRNA-seq platforms is that the cost can limit the complexity of experimental design and therefore the utility of the approach. In this manuscript, we report the use of oligonucleotide barcoding to perform multiplexed CHO cell scRNA-seq to study the impact of tunicamycin (TM), an inducer of the unfolded protein response (UPR). For this experiment, we treated a CHO-K1 GS cell line with 10μg/ml tunicamycin and acquired samples at 1, 2, 4 and 8 hr post-treatment as well as a non-treated TM-control. We transfected cells with sample-specific polyadenylated ssDNA oligonucleotide barcodes enabling us to pool all cells for scRNA-seq. The sample from which each cell originated was subsequently determined by the oligonucleotide barcode sequence. Visualisation of the transcriptome data in a reduced dimensional space confirmed that cells were not only separable by sample but were also distributed according to time post-treatment. These data were subsequently utilised to perform weighted gene co-expression analysis (WGCNA) and uncovered groups of genes associated with TM treatment. For example, the expression of one group of coexpressed genes was found to increase over the time course and were enriched for biological processes associated with ER stress. The use of multiplexed single cell RNA-seq has the potential to reduce the cost associated with higher sample numbers and avoid batch effects for future studies of CHO cell biology.

**Highlights:**

- Polyadenylated ssDNA oligonucleotide labelling is a viable strategy for multiplexed CHO cell scRNA-seq analysis.
- To demonstrate the effectiveness of the method we conducted an experiment to study the CHO cell response to tunicamycin treatment.
- scRNA-seq was carried out on an untreated control and at 4 time points post tunicamycin treatment. Cells from each sample were transfected with a unique oligonucleotide barcode and pooled for single cell transcriptomics.
- Each sample was demultiplexed post-sequencing and gene expression profiles of > 5,300 cells were obtained across the experiment. Following dimensionality reduction and visualisation, the cells were distributed according to sample identity.
- Analysis of the resulting data enabled improved understanding of the transcriptional response to tunicamycin treatment. Three gene coexpression modules were found to be correlated with the tunicamycin time course. Gene set enrichment analysis revealed the over representation of genes related to biological processes associated with ER stress, and protein misfolding in one of these groups of coexpressed genes.
- Further use of this approach will enable the CHO cell biology community to perform increasingly complex single cell experiments in a cost-effective manner.

## 1. Introduction

Chinese hamster ovary (CHO) cells are the most commonly used expression host for the manufacture of recombinant therapeutic proteins such as monoclonal antibodies (Walsh, 2018). Our ability to understand CHO cell biology has improved considerably over the last decade, and the principles of systems biotechnology are being applied to improve the efficiency of CHO cell factories (Kildegaard et al., 2013). Bulk RNA-seq has proven to be an important tool for the analysis of gene expression patterns associated with bioprocess phenotypes (Clarke et al., 2019; Sha et al., 2018). Our laboratory recently demonstrated that scRNA-seq is a powerful method for examining transcriptional heterogeneity across thousands of individual CHO cells (Tzani et al., 2021). Further scRNA-seq analyses have the potential to improve our understanding of how biological heterogeneity affects the performance of a cell culture process as well as the quantity and quality of a recombinant protein produced.

Typical cell isolation protocols for commercially available scRNA-seq platforms require that a single sample is loaded onto an individual microfluidic lane or nanowell cartridge. For studies with large numbers of samples and/or complex experimental designs, the cost can become prohibitive, or batch effects can arise for multiple independent runs. To address these issues, scRNA-seq multiplexing strategies utilising polyadenylated ssDNA containing a sample-specific DNA sequence (barcode) have recently been developed (Cheng et al., 2021). A variety of delivery approaches such as conjugation to an antibody targeting a ubiquitous cell surface protein (Stoeckius et al., 2018), lipid-modification to facilitate attachment of the DNA barcode to the cell membrane (McGinnis et al., 2019) or transfection (Shin et al., 2019) can be used. Once barcoded cells from each sample are pooled for scRNA-seq, the polyA tail of the oligonucleotide allows parallel capture of the barcode and cellular RNA for library preparation. The identity of the sample of origin is determined post-sequencing based on the barcode oligonucleotide associated with each cell.

In this study, we demonstrate that transfected oligonucleotide barcodes allow sample multiplexing for CHO cell scRNA-seq. To illustrate the utility of this approach, we performed an experiment to monitor changes in the CHO cell transcriptome in response to endoplasmic reticulum (ER) stress over an 8 hr time course at single cell resolution.

## 2. Results and discussion

Tunicamycin (TM) inhibits GlcNAc-1-P-transferase, resulting in the disruption of the first step of N-linked protein glycosylation in the ER leading to extensive protein misfolding, ER stress and subsequent activation of the unfolded protein response (UPR) (Brandish et al., 1996; Heifetz et al., 1979; Keller et al., 1979). In this study, a non-producing CHO-K1 GS cell line was treated with 10μg/ml TM to activate the UPR. To confirm that TM induced the ER stress response in the CHO-K1 GS cell line we measured the expression of UPR associated genes *Hspa5, Atf4* and *Ddit3* (Hetz, 2012) as well as the spliced form of *Xbp1*(*i.e., sXBP1*) (Yoshida et al., 2001) using qPCR (Table S1a). For this analysis, samples were obtained at 1, 2, 4, and 8 hr post TM treatment along with an untreated control (TM-) sample. As expected, the expression of each gene increased in each of the TM treated samples over the treatment time course, confirming activation of the UPR in the CHO-K1 GS cell line (Figure 1a; Table S1b).

**Figure 1:**
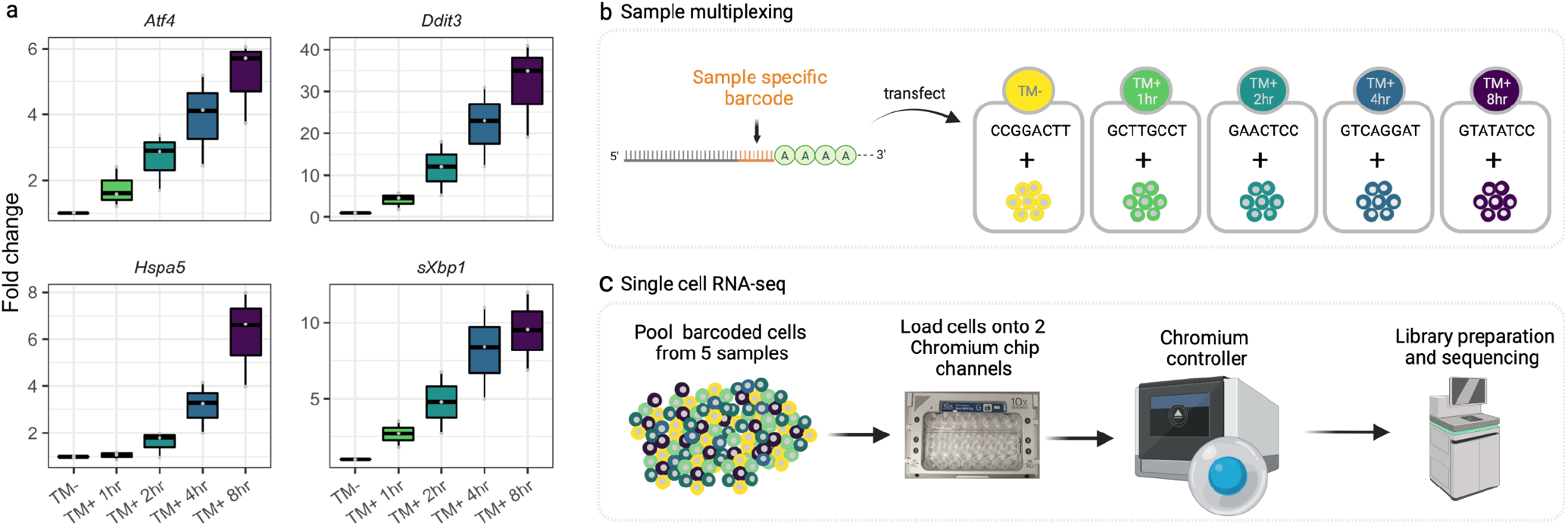
Profiling the temporal CHO cell response to tunicamycin treatment using multiplexed scRNA-seq. A non-producing CHOK1-GS cell line was treated with 10 μg/ml tunicamycin and monitored at selected time points for 8 hr. The **(a)** the upregulation of *Atf4, Ddit3, Hspa5* and *sXbp1* in comparison to the non-TM treated control sample confirmed the induction of the UPR. To prepare samples for multiplex scRNA-seq, **(b)** CHOK1-GS cells were transfected with a short barcode oligonucleotide (SBO) comprising of a sample-specific 8 nt sequence and 30 nt polyadenylated sequence (permitting capture of SBOs in the standard Chromium scRNA-seq protocol). The **(c)** barcoded cells were then pooled and loaded onto two Chromium chip channels. Following cell isolation and size selection of the cellular RNA and SBO cDNA, sequencing libraries from the two GEM wells were prepared. The resulting libraries (2 × cellular RNA and 2 × SBO) were sequenced on an Illumina NovaSeq configured to yield 28 × 91 bp reads.

To demonstrate the effectiveness of multiplexed scRNA-seq for applications in CHO cell biology, we utilised the short barcode oligonucleotide (SBO) method (Shin et al., 2019) modified for compatibility with the 10X Genomics Chromium system. Cells from the TM-sample and each of the 4 time points post-TM treatment were transfected with a SBO containing a unique 8 nt barcode (Figure 1b and Table S2a). Cells from each sample were then pooled and loaded onto two Chromium Chip channels (Figure 1c). During isolation each cell is encapsulated in a Gel Bead-in Emulsion (GEM) droplet in the presence of a bead labelled with sequencing adapters, a cell barcode, unique molecular indexes and a 30bp oligo dT (Zheng et al., 2017). Upon cell lysis, the polyadenylated SBOs are captured and assigned a cell barcode and unique molecular index (UMI) in the same manner as cellular RNA. The preparation of the sequencing libraries from each of the 2 lanes was carried out as described in Chromium Next GEM Single Cell 3’ v3.1 protocol (10X Genomics) until cDNA amplification. At this stage, an SBO additive primer (Table S2b), which recognises the SBO amplification handle, was included in the reaction to increase the SBO product yield. After cDNA amplification the SBO cDNA libraries were separated from the cellular cDNA libraries via the SPRI bead-based clean-up step. The resulting 4 sequencing libraries (2 × cellular RNA and 2 × SBOs) were sequenced using an Illumina NovaSeq 6000 configured to yield 28 × 91 paired-end reads.

The Cell Ranger mkfastq program was used to convert the raw Illumina BCL base call files to FASTQ files containing the sequencing reads for the two RNA and two SBO libraries (1 from each GEM well). To generate a reference index we merged the Chinese hamster PICR nuclear genome (Rupp et al., 2018) with the mitochondrial genome of the CHO-K1 cell line (Xu et al., 2011). Kallisto | bustools (Melsted et al., 2021) was used to pseudoalign reads from the RNA libraries and generate UMI counts for each cellular barcode (Figure S1a). The 2 raw UMI count matrices generated by Kallisto | bustools were imported into the R statistical software environment and empty droplets were removed using the knee plot method (Macosko et al., 2015) (Figure S2a & S2b) and Seurat (Hao et al., 2021) objects were created. miQC (Hippen et al., 2021) was used to capture the relationship between the %UMIs from mtDNA genes and the total number of genes detected and subsequently eliminate low quality cell barcodes which likely are from cells with damaged membranes (Luecken and Theis, 2019). The remaining cell barcodes were removed from further analysis if the number of UMIs and/or the number of genes detected were 3 median absolute deviations (MAD) from the median of the population. In total, 3,201 and 3,381 high quality cells were retained from the RNA 1 and RNA 2 libraries respectively (Figure S3a & S3b).

To integrate the SBO data with their matched RNA library, CITE-seq count (Roelli et al., 2019) (Figure S1b) was used to generate SBO-UMI count matrices. Only those cell labels which were present in both the RNA and SBO data captured from the same GEM well were retained for further analysis. Next, the SBO UMI count matrices were center-log-ratio (CLR) transformed in Seurat and the HTODemux function was used to determine the sample from which each cell originated. A 0.94 classification threshold for HTODemux was selected (Figure S4) to maximise the number of singlets (cell associated with a single SBO) across both GEM wells (Figure S5). Of the 6,582 cells detected in both the cellular RNA and SBO libraries, 80% (n = 5,314) could be classified as originating from one of the 5 samples (Figure 2a; Table S3a); The overall number of singlet cells identified from GEM well 1 (n = 2,645) and GEM well 2 (n = 2669) was consistent (Figure S6a). The number of singlet cells identified ranged from 1,358 for the TM+2hr sample, to 803 for the TM+ 4hr sample. While the number of negative cells (n = 331) for this sample was significantly higher, the origin of this variation is unclear. Both GEM wells did, however, have a comparable number of negative cells (Figure S6b, Table S3b & S3c) indicating that the variation occurred prior to loading of cells onto the Chromium platform.

**Figure 2:**
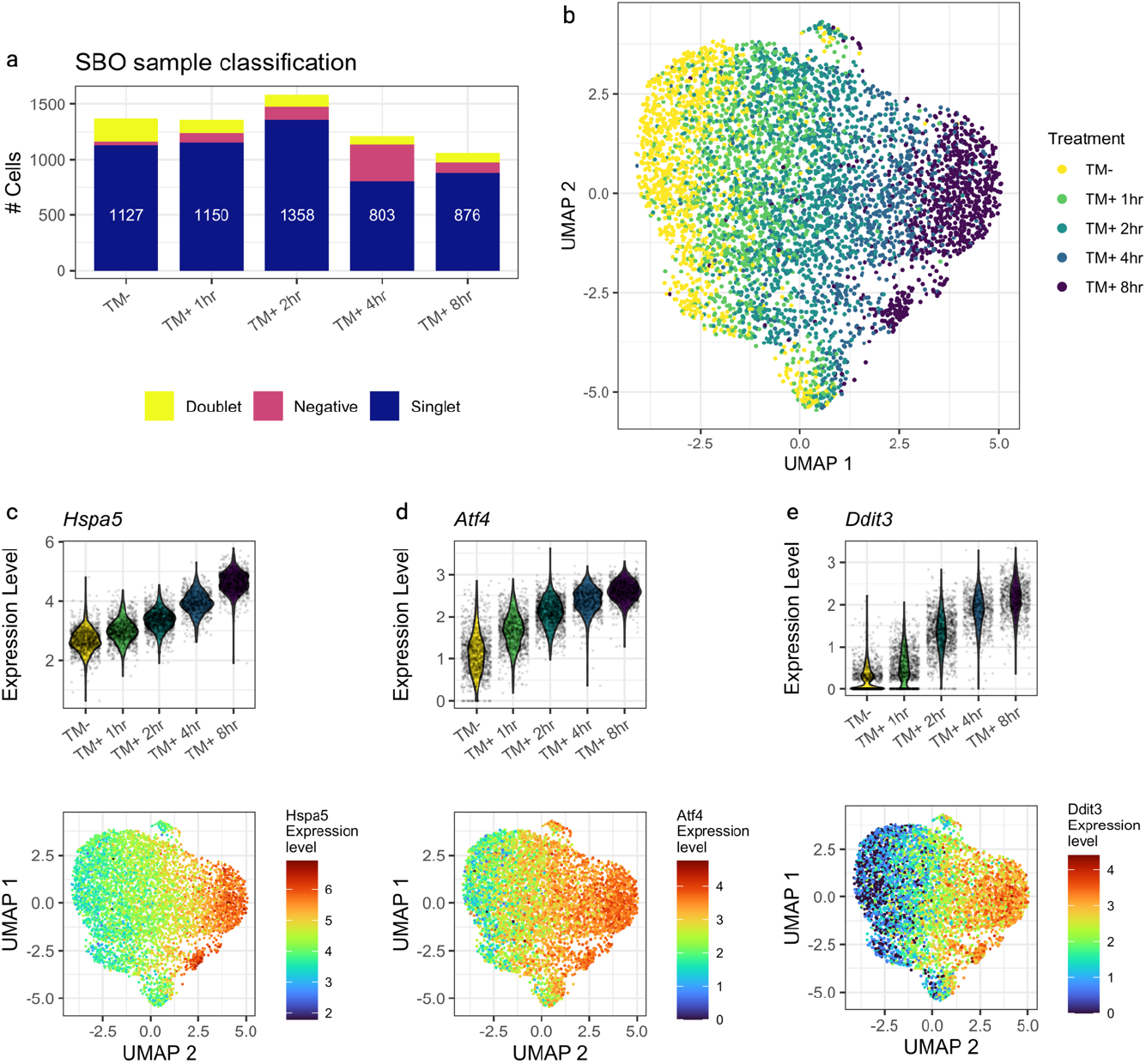
Sample barcode oligonucleotides enable multiplexed CHO cell scRNA-seq. Demultiplexing of the scRNA-seq data via classification of the oligonucleotide labels resulting in **(a)** 5,314 cells confidently assigned to one of the 5 samples. The number of singlet, negative and doublet classifications are provided in Table S3. **(b)** UMAP dimensionality reduction of the gene expression data and overlay of the sample label confirms that the cells distribute according to the TM treatment and timepoint. Analysis of the **(c)** *Hspa5*, **(d)** *Atf4* and **(e)** *Ddit3* gene expression illustrates the concordance of the demultiplexed single cell data to that of the qPCR expression data and clear changes in expression across the UMAP space.

The demultiplexed gene expression data for GEM well 1 and GEM well 2 were merged into a single Seurat object for further analysis. The SCTransform algorithm (Hafemeister and Satija, 2019) was used to normalise the UMI count data, identify the 3,000 most variable genes and remove variation arising in the number of genes detected per cell. The cell cycle phase of each cell was also estimated (Figure S7) and regressed out. Next, to obtain a global overview distribution of cells in each sample, multidimensional visualisation was carried out using uniform manifold approximation and projection (UMAP) on 20 principal components (Figure S8). From the resulting plots, we confirmed that there was no significant batch effect between the gene expression acquired from the 2 GEM wells (Figure S9a) and that the effect of cell cycle had been successfully removed (Figure S9b). Labelling each cell by the treatment condition illustrates that the cells are distributed in the 2-dimensional UMAP plot according to the treatment and timepoint indicating that demultiplexing of samples using the SBO method was successful (Figure 2b). Furthermore, the expression of *Hspa5* (Figure 2c), *Atf4* (Figure 2d) and *Ddit3* (Figure 2e) observed with scRNA-seq were comparable to that of the qPCR analysis (Figure 1a).

To further understand the CHO cell transcriptional response to TM treatment, we performed weighted gene coexpression network analysis (WGCNA) for scRNA-seq (Langfelder and Horvath, 2008; Morabito et al., 2021). WGCNA is a widely used technique that utilises a “guilt by association” based method to elucidate groups of genes that are coexpressed genes (i.e., modules). While WGCNA has been widely used for bulk gene expression data captured from microarray and RNA-seq, the inherent sparsity of single cell RNA-seq data presents a challenge for WGCNA. To overcome the issue of sparsity, we utilised a previously developed method to create pseudobulk profiles (termed metacells) from the average of *k* nearest neighbour cells in the UMAP space (Morabito et al., 2021). Here, we selected the 3,000 highly variable genes identified by SCTransform and aggregated cells separately for each sample in our experiment. The average expression was determined for each dataset by considering 60 nearest neighbour cells yielding a total of 2,159 metacells for each of the 5 samples (Figure S10a & S10b).

The metacell expression matrix was then utilised to perform WGCNA with power (β) = 7 and the minimum number of genes per module = 50 (see methods for further details) were selected to approximate a scale free topology (Figure S11), resulting in the identification of 9 coexpression modules (each module was assigned an arbitrary colour for reference) ranging from 483 to 60 genes were identified (Figure 3a & 3b; Table S4). A measure, known as the module eigengene (ME) (Zhang and Horvath, 2005), was used to summarise the dominant expression pattern across the genes in each of 9 modules identified by WGCNA. The ME is calculated by performing principal components analysis on the metacell data for genes within a particular module. The values in 9 MEs were separated based on sample and plotted to assess the correlation of each module with the transcriptional response to TM treatment (Figure S12). The intramodular connectivity (kME) was also calculated by correlating the expression of each gene to the MEs to enable the identification of hub genes (i.e., those genes which are most similar to that of the overall expression of their parent module) (Table S4).

**Figure 3:**
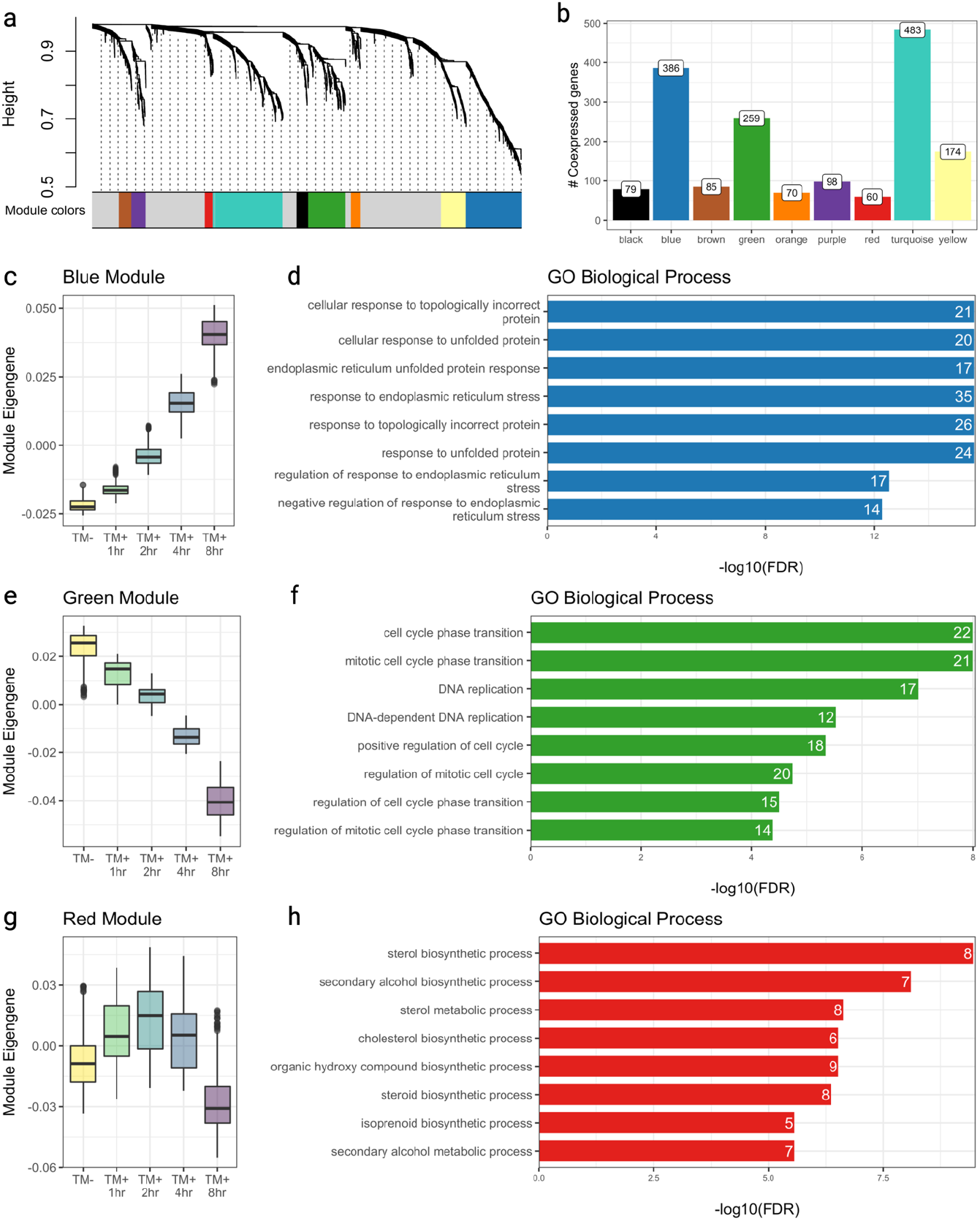
Identification of coexpressed gene modules associated with the temporal response to tunicamycin treatment. WGCNA identified **(a)** 9 coexpressed modules ranging from **(b)** 483 to 60 genes. To determine if the identified groups were associated with the tunicamycin treatment time course, the expression data were stratified for each module and principal components analysis was carried out to obtain the module eigengene (i.e., the first principal component). The **(c)** blue module eigengene expression indicated that these genes tended to increase post-TM treatment. Following gene set enrichment analysis against GO biological process categories related to **(d)** ER stress and protein folding were found to be significantly overrepresented in this module. In contrast, green module genes tended to **(e)** decrease in expression following TM treatment and genes related to cell cycle were found to be **(f**) enriched in this group of genes. Genes were found to be altered with tunicamycin treatment and time. Red module genes tended to **(g)** increase in expression at 1hr and 2hr post TM treatment in comparison to the TM-control, before decreasing at the 4 hr and 8hr time points. The genes in this module were **(h)** enriched for categories including sterol and cholesterol metabolism.

Following examination of the MEs, the blue, green, and red MEs were found to be associated with TM treatment and were prioritised for further analysis. For the blue coexpression module, the ME was found to increase with treatment duration (Figure 3c) indicating that the 386 coexpressed genes in this module tended to increase in expression following TM treatment. Examination of the blue kME values revealed genes such as *Hspa5* were amongst the top ranked hub genes in this module (Figure S13; Table S4). Categories related to ER stress, and the unfolded protein response were overrepresented within the blue module following gene set enrichment analysis against GO biological processes (Figure 3d; Table S5a). In contrast, the green ME genes decreased following TM treatment (Figure 3e, Figure S14, Table S4b) and following enrichment analysis we found that biological processes relating to cell cycle processes (e.g., cell cycle transition) were overexpressed within the 259 genes in this module (Figure 3f; and Table S5b). This result is in agreement with previous reports in the literature showing that tunicamycin affects cell cycle (Brewer et al., 1999; Wang et al., 2011). Indeed, from the cycle classification carried out in this study (Figure S7), we observed an increase in the cells predicted to be in the G1 phase and a decrease of cells in S phase over the 8 hrs post TM treatment. In the case of the red module (Figure S15, Table S4c), the ME increased at 1 hr and 2hrs post TM treatment in comparison to the control sample posttreatment followed by a decrease at 4 hrs and 8hrs (Figure 3g). Despite the red module containing the lowest number of coexpressed genes, we however observed the significant enrichment of several metabolic processes including cholesterol metabolism (Figure 3h; Table S5c). This finding is consistent with a previous report showing that during ER stress changes in the expression of some genes (e.g. *Hmgcr*) involved in lipogenesis/cholesterol biosynthesis can be initially upregulated, peak at 2-4 hrs and are then reduced by 8 hrs (Arensdorf et al., 2013).

## Conclusion

In this study, we have demonstrated the effectiveness of transfecting cells with polyadenylated ssDNA oligonucleotide barcode for scRNA-seq sample multiplexing. The technique enabled cells originating from multiple samples to be pooled and loaded onto the same Chromium Chip channel, significantly reducing cost per sample. Multiplexed scRNA-seq will enable more complex experimental designs to be utilised to enhance the study of CHO cell biology using this technology in the future.

## 3. Materials and Methods

### 3.1 CHO cell culture and tunicamycin treatment

A non-producing CHOK1-GS cell line was seeded at a density of 2 × 10^5^ cells/ml in 5ml CD FortiCHO™ medium (Gibco, cat.no. A1148301) supplemented with 4mM L-glutamine (L-Glutamine, cat.no. 25030024) in bioreactor tubes (Corning™ cat. no 431720). The cultures were maintained at 37°C, 170 rpm, 5% CO2 and 80% humidity in a shaking incubator (Kuhner). The cells were treated with 10μg/ml tunicamycin (Cell Signalling Technology®, cat.no.12819S) at 48 hr post-seeding and samples taken at 1 hr, 2 hr, 4 hr and 8 hr. An untreated control sample was also obtained.

### 3.2 qPCR

Total RNA was extracted from cell pellets using Trizol (Invitrogen, cat. no. 15596026). The purity and quantity of RNA were assessed by spectrophotometry (Nanodrop 1000; Thermo Fisher Scientific). cDNA was generated from 2 μg of total RNA using the Applied Biosystems kit (ref 4368814), according to the manufacturer’s guidelines. qPCR was performed on a QuantStudio™ 3 Real-Time PCR System (Applied Biosystems™) in 20 μl reactions with the Fast SYBR™ Green Master Mix (Applied Biosystems™, cat.no.4385616), primers (Table S1a) at 0.4μM final concentration and 4 μl of 1:10 diluted cDNA. Reactions minus reverse transcriptase were included to control for contaminating genomic DNA. RNA levels of the measured ER stress markers (*Atf4, Hspa5, Ddit3* and *sXBP1*, Table S1b) from tunicamycin treated samples were normalised to *Gusb* RNA levels. Relative RNA quantification calculated using the Livak method (2^−ΔΔCT^) (Livak and Schmittgen, 2001) was used to compare each TM+ sample to the TM-control (Table S1b).

### 3.3 Single cell RNA-seq

#### 3.3.1 SBO transfection and sample pooling

Transfection of CHO cells with sample specific SBOs (Table S2a) (42nM) with of 22 ug/ml PEI was performed for all timepoints 1 hour before to sample collection (i.e., the TM+ 8hr, 4hr, 2hr samples were transfected at 7-, 3- and 1-hour post TM treatment respectively while the TM+ 1 hr sample, tunicamycin treatment and transfection was performed simultaneously). Cells were counted and pooled following the cell sample preparation guide from 10X Genomics (CG00053 • Rev C). Briefly, cells were centrifuged at 200 rcf for 3 minutes, the supernatant was removed, and the cells were resuspended in PBS with 0.04% BSA with gentle mixing. Cells were passed through a 35 μM strainer (Falcon™ cat. No. 352235) to remove cell debris and large clumps. Cell concentration was determined with the Luna™ automated cell counter (Thermo Fisher Scientific) and equal number of cells from the different treatment conditions were pooled. Following sample pooling, the cell concentration was measured again (total conc. = 1,180 cells/μl, viability = 94.3%) before to loading onto two channels of a Chromium Chip.

#### 3.3.2 Library preparation

For the generation of cDNA and the matched SBO libraries (1 per GEM well), the 10X Genomics Single cell 3’ v3.1 workflow (CG000204 • Rev D) was followed until the cDNA amplification step. At this stage the SBO additive primer (Table S2b) was added to the cDNA amplification reaction at 0.015μM final concentration. Following cDNA amplification, a clean-up step with 0.6X SPRI beads (Beckman Coulter™, B23317) was performed according to the 10X Genomics protocol. The SBO-derived cDNAs are shorter than the mRNA-derived cDNAs and were therefore present in the supernatant and the beads respectively of the clean-up step. The libraries derived from cellular mRNAs were prepared following the 10X Genomics Single cell 3’ v3.1 protocol (GC000204 • Rev D).

The supernatant from the 0.6X SPRI clean-up step following cDNA amplification, which contained the SBO-derived cDNA was transferred to a new tube and was further purified with two 2X SPRI purification steps. Briefly, 1.4X volume of SPRI beads was added to the SBO-derived cDNA containing solution and following an 80% ethanol wash and elution step, an additional round of 2X SPRI beads purification was performed. The eluate was used for the SBO library preparation using a previously described protocol (Shin et al., 2019). The primers (Table S2b) used for the 1st PCR for the amplification of the SBO libraries were: SBO-PCR-F and SBO-PCR1-R (3 min at 95°C; eight cycles of 20 s at 95°C, 20 s at 64°C, 20 sec at 72°C; 5 min at 72°C). For the second PCR the same forward primer (SBO-PCR-F) and two reverse primers SBO-PCR2-R1 or SBO-PCR2-R2 were used to differentiate (3min at 95°C; six cycles of 20 s at 95°C, 20 s at 60°C, 20 s at 72°C;5 min at 72°C) the samples loaded onto the two different Chromium chip channels. The first PCR product was purified with a 1.8X SPRI bead purification step while the second PCR product was purified by size selection from an 8% acrylamide gel.

Prior to sequencing concentration of all libraries was measured with the KAPA Library Quantification kit (KK4824, Roche) following manufacturer’s guidelines and library size was assessed with TapeStation (Agilent) traces.

#### 3.3.3 Sequencing

The cellular RNA and SBO libraries were pooled and loaded onto an SP flowcell of an Illumina NovaSeq configured to yield 28 × 91 bp paired-end reads. The mkfastq programme (10x Genomics Cell Ranger v6.0.2) was utilised to convert the resulting Illumina BCL files to FASTQ for further analysis.

### 3.4 scRNA-seq data analysis

#### 3.4.1 scRNA-seq read mapping and quality control

A hybrid reference genome comprised of the CriGri-PICR nuclear genome (GCA_003668045.1) (Rupp et al., 2018) and mitochondrial DNA sequence of the CHO-K1 cell line (GCA_000223135.1) (Xu et al., 2011) was constructed for scRNA-seq mapping. The cellular RNA libraries from the 2 GEM wells were separately pseudoaligned to the reference and counted using Kallisto | bustools (Melsted et al., 2021) (Figure S1). The RNA count matrices were imported into R and empty droplets were removed using the knee plot method (Macosko et al., 2015) and a cell barcode whitelist was produced for CITE-seq count (see Section 3.4.2). Seurat v4 (Hao et al., 2021) objects were then created for each GEM well library and low quality cell barcodes were removed by first assessing the relationship between the number of UMIs originating from mtDNA and the number of detected genes in each cell using miQC (Hippen et al., 2021). In addition, cell barcodes where the number of genes detected and/or number of UMIs were 3 median absolute deviations (MAD) outside the median of each population were eliminated.

#### 3.4.2 Analysis of SBO libraries

The FASTQ files for the SBO libraries from the 2 GEM wells were first processed individually using CITE-Seq count. The whitelist of cell barcodes generated from the Kallisto | bustools kb-count and following the filtering of empty droplets using the knee plot. CITE-Seq count was configured to count the 8nt unique sample barcode for each cell barcode via the following options: -start-trim 22; -cbf 1; - cbl 16; -umif 17; -umil 28; --bc_collapsing_dist 1; --max-error 1; --sliding-window; -cells 5000. The resulting count matrix for the two SBO libraries was imported into R.

#### 3.4.3 Sample demultiplexing

Only those cell barcodes present in both the cellular RNA and matched SBO library were retained for further analysis. The SBO counts generated by CITE-seq were added as a new assay to the cellular RNA Seurat objects from each GEM well. before the HTODemux function implemented in Seurat (positive.quantile = 0.94) was used to classify the sample of origin for each cell. Negative classifications or where two SBOs were associated with one cell (i.e., doublets) were removed from further analysis. The two Seurat objects with sample classifications were then merged for further analysis (Figure S1).

#### 3.4.3 scRNA-seq data normalisation and visualisation

Cellular RNA-seq data were normalised by SCTransform (Hafemeister and Satija, 2019) and variability due to the number of genes detected per cell was removed. We also mapped the precompiled S phase and G2M phase gene lists provided in Seurat to Ensembl Chinese hamster gene IDs and used the CellCycleScoring Seurat function to classify cells in each phase and to regress out variability due to cell cycle. The normalised data for 3,000 variable genes identified by SCTransform were used to conduct PCA and the first 20 principal components were used for UMAP visualisation of the scRNA-seq data in a reduced dimensional space.

#### 3.4.4 Weighted gene coexpression network analysis

The scWGCNA (Morabito et al., 2021) and WGCNA R (Langfelder and Horvath, 2008) packages were utilised to perform coexpression analysis. The demultiplexed, normalised data for the 3,000 variable genes were first used to generate pseudobulk “metacells” (to alleviate scRNA-seq sparsity) by merging the expression of the 60 nearest neighbours of each cell. The aggregation process was performed separately for the cells in each of the 5 samples (TM-, TM 1hr, TM 2hr, TM 4hr and TM 8hr) and then combined into a single metacell expression matrix for WGCNA. Next, the bi-weighted mid-correlation between all pair of genes was calculated to construct a signed similarity matrix. The similarity matrix was raised to a series of powers (β), to determine the agreement of each transformed matrix with a network with scale free topology (SFT). WGCNA requires only an approximate fit to an ideal SFT (Zhang and Horvath, 2005). The first β that resulted in a SFT R^2^ > 0.8 and mean connectivity < 100 was selected to create the adjacency matrix. A specialised network distance metric termed the topological overlap measure (TOM) was then used to determine the coexpression similarity between pairs of genes in the adjacency matrix. The TOM assesses not only the direct correlation between genes but also the degree of agreement between the genes and their relationship with all other genes. The WGCNA blockwiseConsensusModules function was used to identify groups of coexpressed genes (i.e., modules).

An expression summary measure was calculated for each coexpression module by sub-setting the metacell expression matrix for each group of coexpressed genes and the performing PCA. The first principal component, referred to as the module eigengene (ME), was retained and the values were plotted for each sample to determine if there was an association between groups of genes and tunicamycin treatment. In addition, a measure termed the intra-modular connectivity (kME) was also calculated by determining the correlation of individual genes to the MEs to identify hub genes within each coexpression module.

#### 3.4.4 Enrichment Analysis

The overrepresentation of gene ontology (GO) biological processes in coexpressed gene modules were assessed with the R WebGestaltR package (Wang et al., 2017). Enriched biological processes with a Benjamini-Hochberg adjusted p-value of < 0.05 were considered significant.

## Abbreviations

CLR: center-log-ratio
CHO: Chinese hamster ovary
ER: Endoplasmic reticulum
GO: Gene Ontology
MAD: Median Absolute Deviations
ME: Module Eigengene
scRNA-seq: single cell RNA-seq
SBO: short barcode oligonucleotide
SFT: Scale Free Topology
TOM: Topological Overlap Measure
TM: Tunicamycin
t-SNE: t-stochastic neighbour embedding
UPR: unfolded protein response
UMAP: uniform manifold approximation and projection
WGCNA: weighted gene coexpression network analysis
ssDNA: single stranded DNA

## Data availability

The scRNA-seq data have been deposited to the Sequence Read Archive (SRA) (accession: PRJNA821033). The code required to reproduce the results presented in this manuscript are available at: https://clarke-lab.github.io/multiplexed_CHO_scrnaseq

## Author Contributions

IT and CC conceived the study and designed experiments; Cell culture and scRNA-seq were carried by I.T and SB. scRNA-seq data analysis was performed by MCR, and CC. IT, SB, and CC wrote the manuscript. All authors reviewed the paper.

## Acknowledgements

The authors gratefully acknowledge funding from Science Foundation Ireland (grant reference: 15/CDA/3259). Figures were created with BioRender.com.

## Supplementary Tables

**Table S1: Assessment of ER stress marker gene expression using qPCR.** To determine the impact of tunicamycin treatment on the CHO-K1 GS cell line, **(a)** primers were designed to measure the expression of *Hspa5, Ddit3, Atf4* and spliced *Xbp1*. The **(b)** fold change of each of these targets at each selected time-point relative to the non-treated control sample was calculated using the 2^−ΔΔCT^ method.

**Download:** https://app.box.com/s/dm0urmkoo671lwn45yeh3ujqv0nshefs

**Table S2: Sample specific barcoding of samples for scRNA-seq.** Each sample was **(a)** labelled with a polyadenylated ssDNA oligonucleotide with an 8nt barcode. During library preparation a **(b)** series of primers were used to amplify the SBOs for sequencing.

**Download:** https://app.box.com/s/nni24bk1kxql0ndpdcfip50vsyfoqn1a

**Table S3: HTODemux classifications.** The number of singlet, negative and doublet identifications from the SBO data are shown for **(a)** all cells as well as the classifications for **(b)** GEM well 1 and **(c)** GEM well 2.

**Download:** https://app.box.com/s/zydme26a1ccj4ekp9v50ulqbcwug94gn

**Table S4: Identification of coexpression modules using WGCNA.** WGCNA identified 9 coexpression modules from the CHO cell scRNA-seq data. The Ensembl ID, biotype, symbol, and description are shown for each gene along with the intramodular connectivity (kME) for each coexpression module.

**Download:** https://app.box.com/s/x1rw8bsdrn6lur8awssfdwq4chofi2be

**Table S5: Enrichment analysis of coexpression modules associated with TM treatment.** Following the identification of coexpression modules that were correlated with the TM time course, gene set enrichment analysis was carried out to determine if GO biological processes were overrepresented in the **(a)** blue, **(b)** green and **(c)** red modules. GO categories found to be enriched along with the Benjamini-Hochberg adjusted p-value and the genes leading to the enrichment observed are shown.

**Download:** https://app.box.com/s/s3ddeiftxtnc4udjevlakh73kgzr060n

## Supplementary Figures

**Figure S1:**
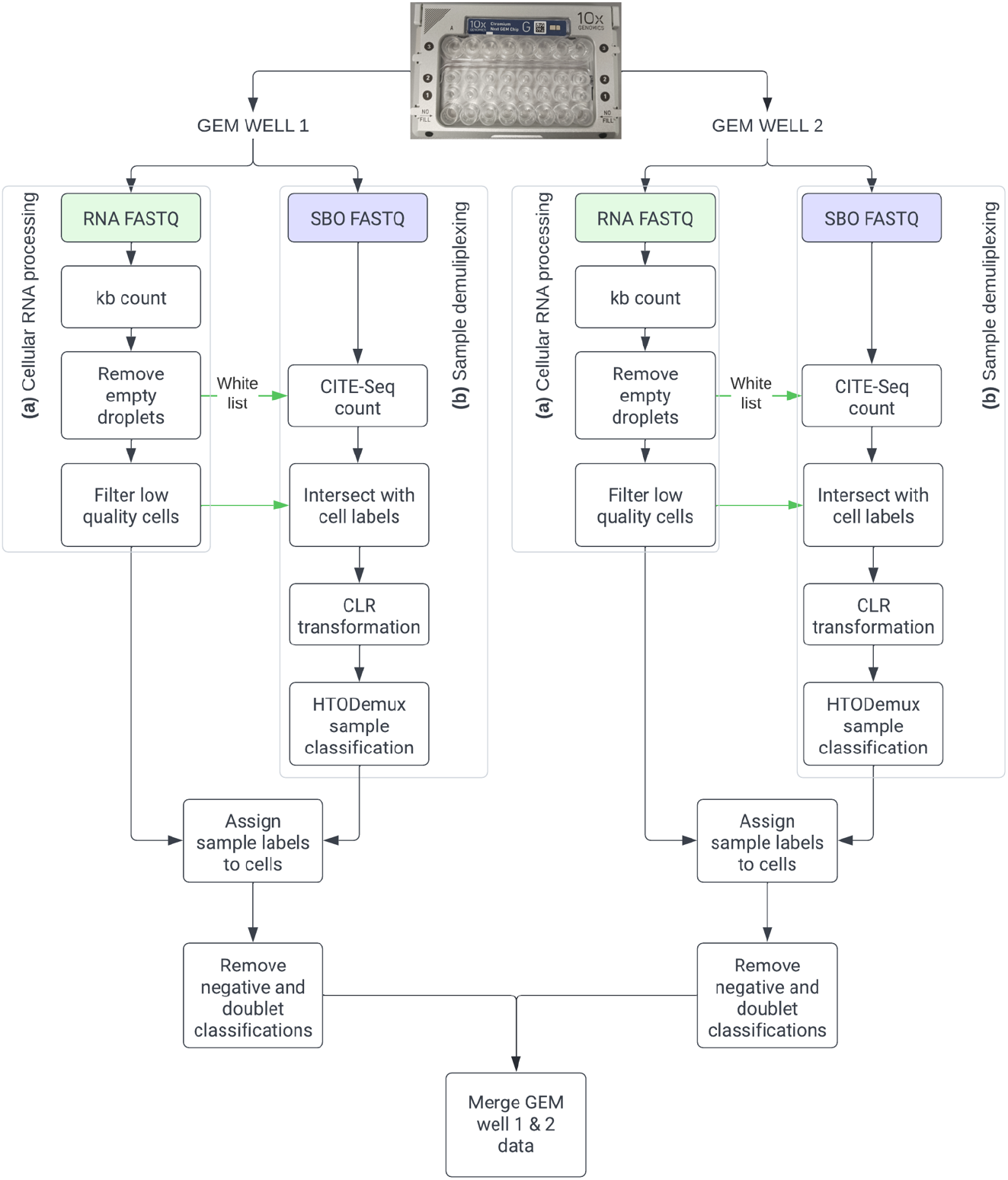
Bioinformatics workflow for scRNA-seq data QC and SBO-based demultiplexing. Aliquots of pooled SBO tagged CHO cells were loaded onto two channels of the Chromium for scRNA-seq analysis. Cellular RNA and SBO libraries were independently prepared from each channel and sequenced. The FASTQ files for **(a)** the cellular RNA data were pseudoaligned against a combined reference index constructed from the Chinese hamster nuclear and CHO-K1 mitochondrial genome using Kallisto | bustools and an initial UMI-gene count matrix generated. Following the removal of empty droplets, the UMI cell-gene count matrix was imported into Seurat and low-quality cells were removed based on the %UMIs from the mitochondria, as well as the number of UMIs and genes detected. The **(b)** SBO FASTQ files were analysed using CITE-seq count to produce a UMI-SBO matrix for non-empty droplets. The cell labels from each of the RNA libraries were intersected with the SBO cell labels from the corresponding GEM well and non-overlapping cell barcodes were discarded. To determine the sample identity, the SBO data was CLR transformed before HTODemux was used to classify each cell’s sample of origin. Those cells classified as negative or doublet were removed before the data from GEM well 1 and GEM well 2 were merged for further analysis (i.e., UMAP visualisation and WGCNA).

**Figure S2:**
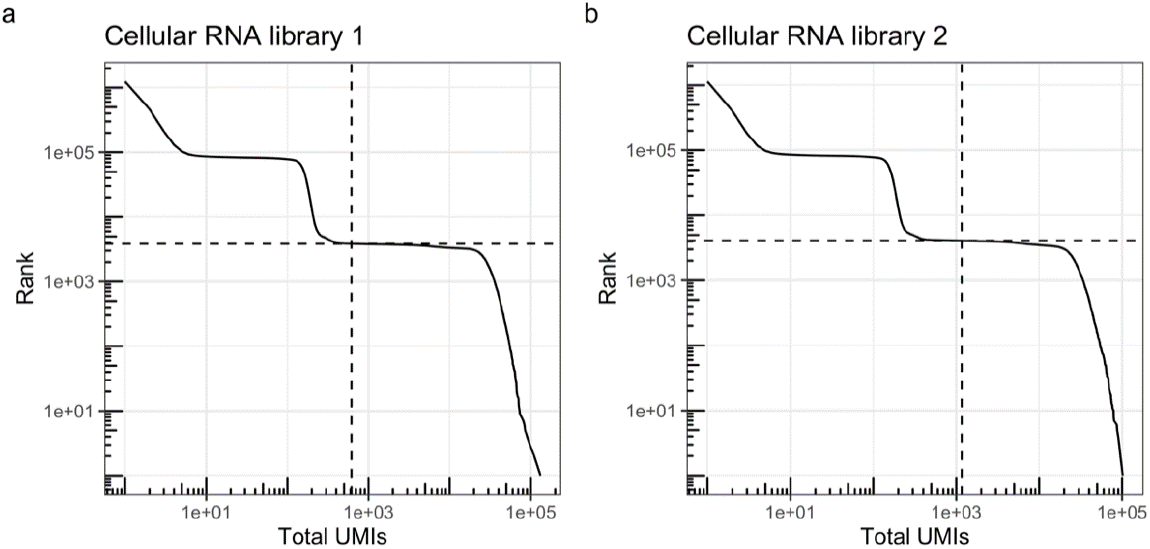
Removal of empty droplets. Knee plots illustrating the removal of empty droplets for the cellular RNA data for **(a)** GEM well 1 and **(b)** GEM well 2 based on ranking the total UMI for counts per cellular barcode.

**Figure S3:**
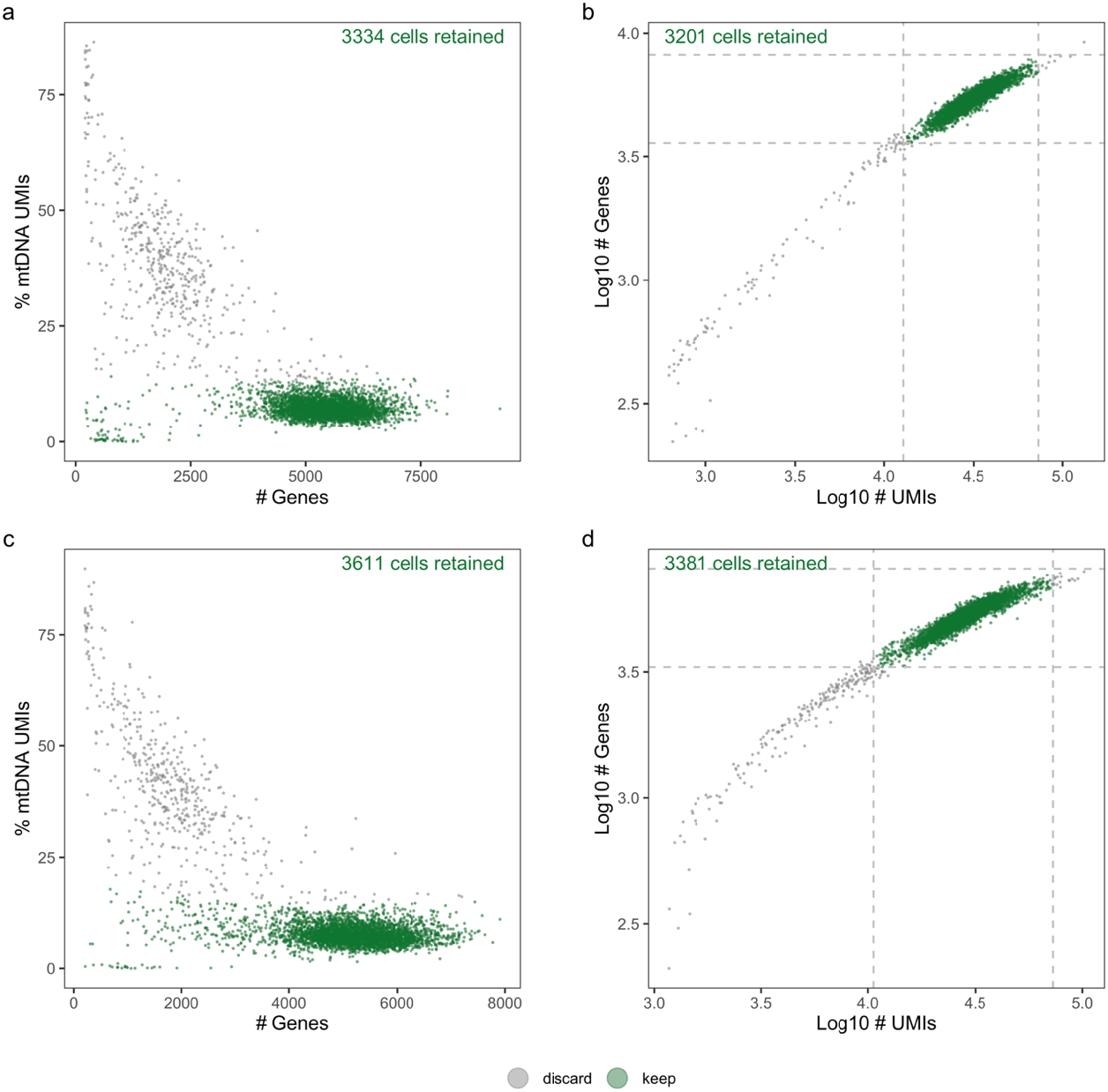
Pre-processing of cell-gene UMI count matrices. miQC was used to assess the relationship between the number of UMIs originating from mtDNA and the number genes detected to identify low quality cells in the cellular RNA libraries from **(a)** GEM well 1 and **(c)** GEM well 2. If the number of UMIs or number of genes detected per cell was beyond 3 MAD of the median of the population of **(b)** GEM well 1 or **(d)** GEM well 2 those cells were also discarded.

**Figure S4:**
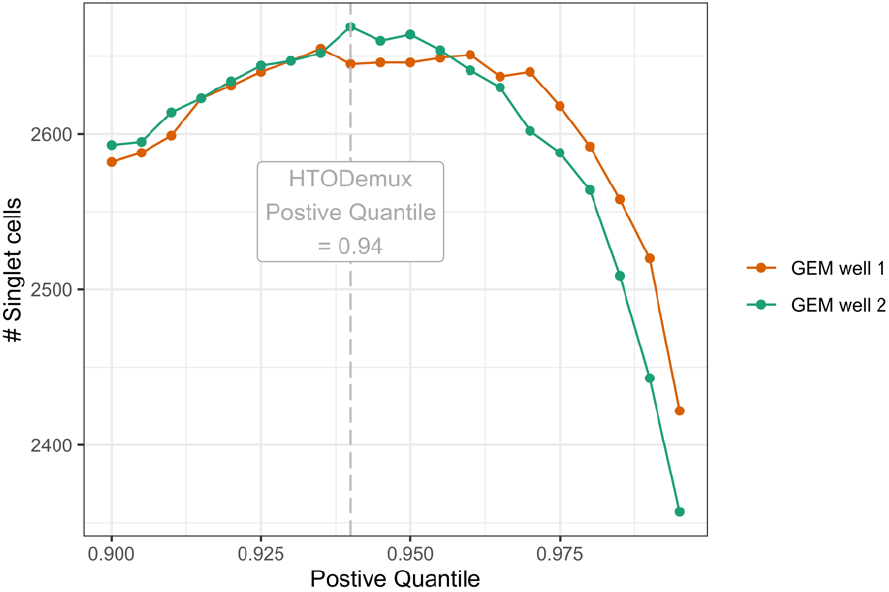
Selection of the HTODemux positive quantile threshold for SBO-based sample classification. A range of positive quantile value from 0.9 to 0.999999 were assessed to identify a value for both GEM well 1 and GEM well 2 datasets. 0.94 was selected as the classification threshold to maximise the number of singlet cells.

**Figure S5:**
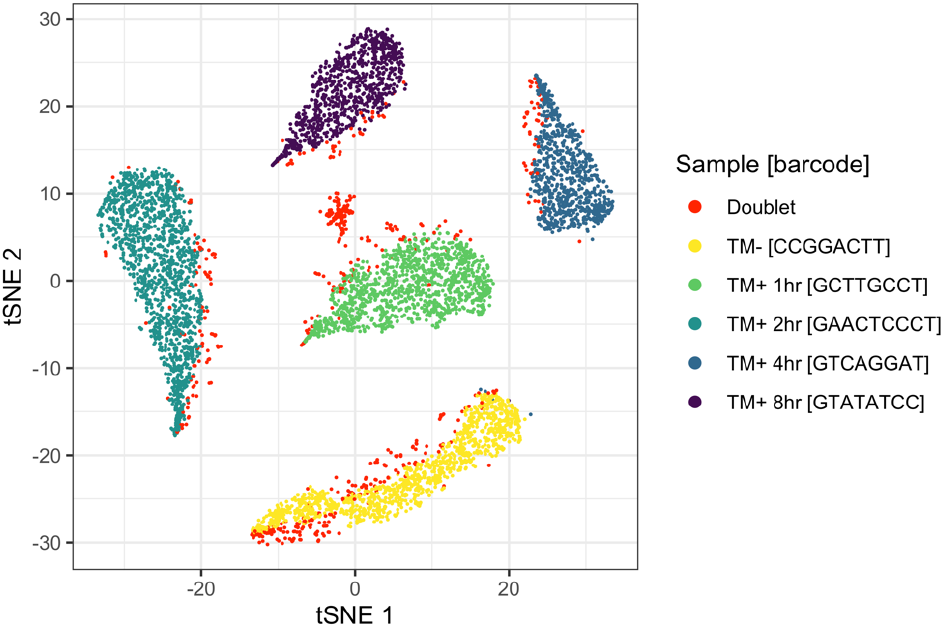
Visualisation of the classified SBO count matrix. t-Stochastic neighbour embedding (tSNE) dimensionality reduction and visualisation of the combined GEM well 1 and GEM well 2 UMI-SBO count matrices. HTODemux classifications are shown along with those cells identified as doublets.

**Figure S6:**
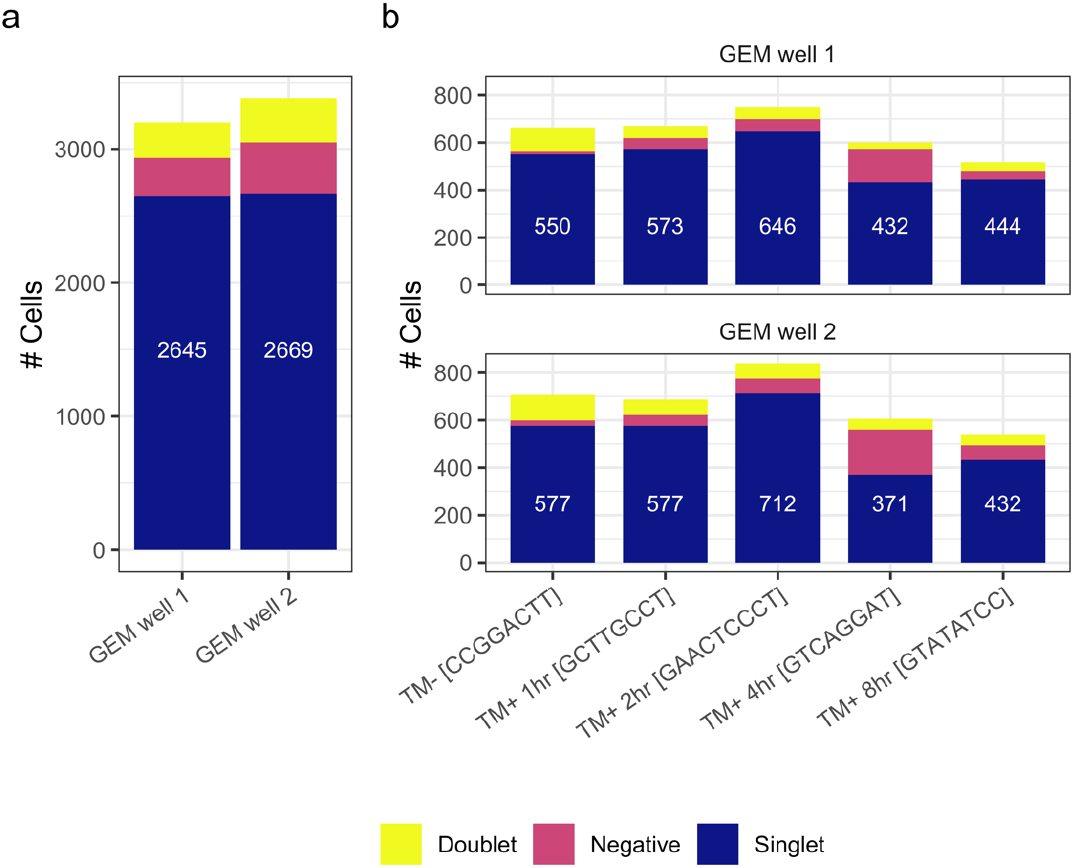
Demultiplexing cells based on SBO labelling. HTODemux with a positive quantile threshold = 0.94 was used to identify a total of **(a)** 2,645 and 2,669 singlets from GEM well 1 and GEM well 2 respectively. The **(b, c)** sample classifications across the 2 GEM wells were comparable. The number of singlet classifications are shown in white; the number of negative and doublet classifications are provided in Table S3. For the TM+ 4hr sample, a comparable number of SBO negative cells was observed for GEM well 1 (n = 141) and GEM well 2 (n = 190). While the origin of this variation is unknown these data indicate that the difference in detection rate of TM+ 4hr arose from the experimental steps conducted prior to cell isolation.

**Figure S7:**
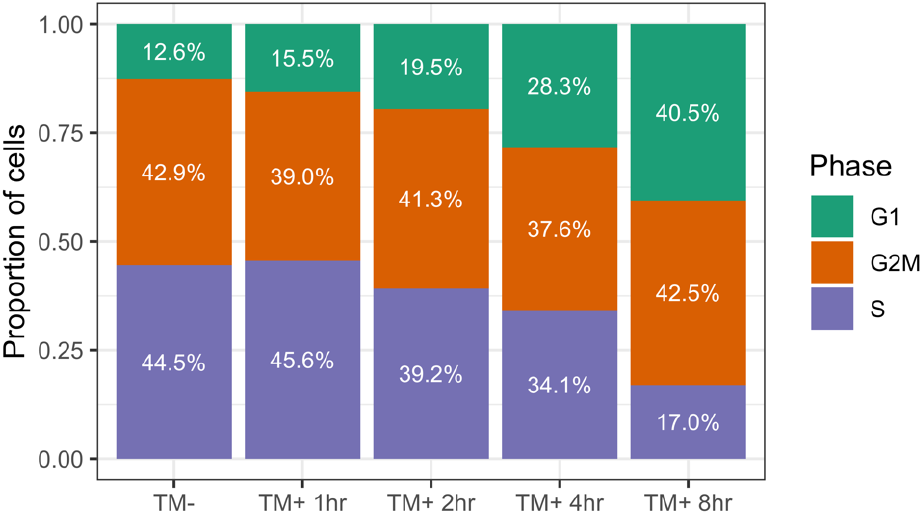
Classification of cell cycle phase. The Seurat CellCycleScoring function was used to classify the phase of cell cycle for the 5,314 cells based on the expression of genes associated with the S phase and the G2M phase (a cell not classified in either phase is designated as G1). The proportion of cells predicted to be in each phase for each of the 5 samples is shown in white. The likelihood of each cell being either S or G2M is also outputted by the algorithm and was used to regress the effect of cell cycle from the data during SCTransform normalisation.

**Figure S8:**
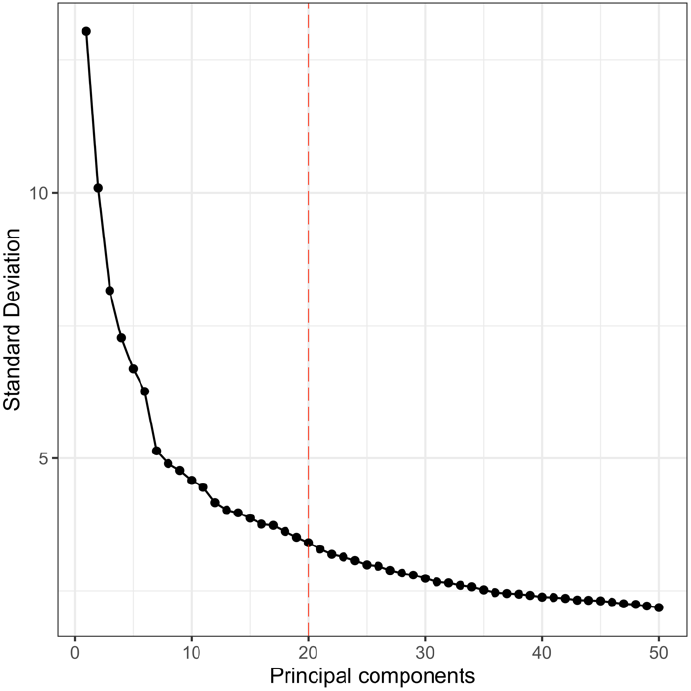
Selection of principal components for UMAP: Principal components analysis was performed on the SCTransform normalised gene expression data (cells = 5,314, genes = 3000). The first 20 principal components were retained for the UMAP visualisation.

**Figure S9:**
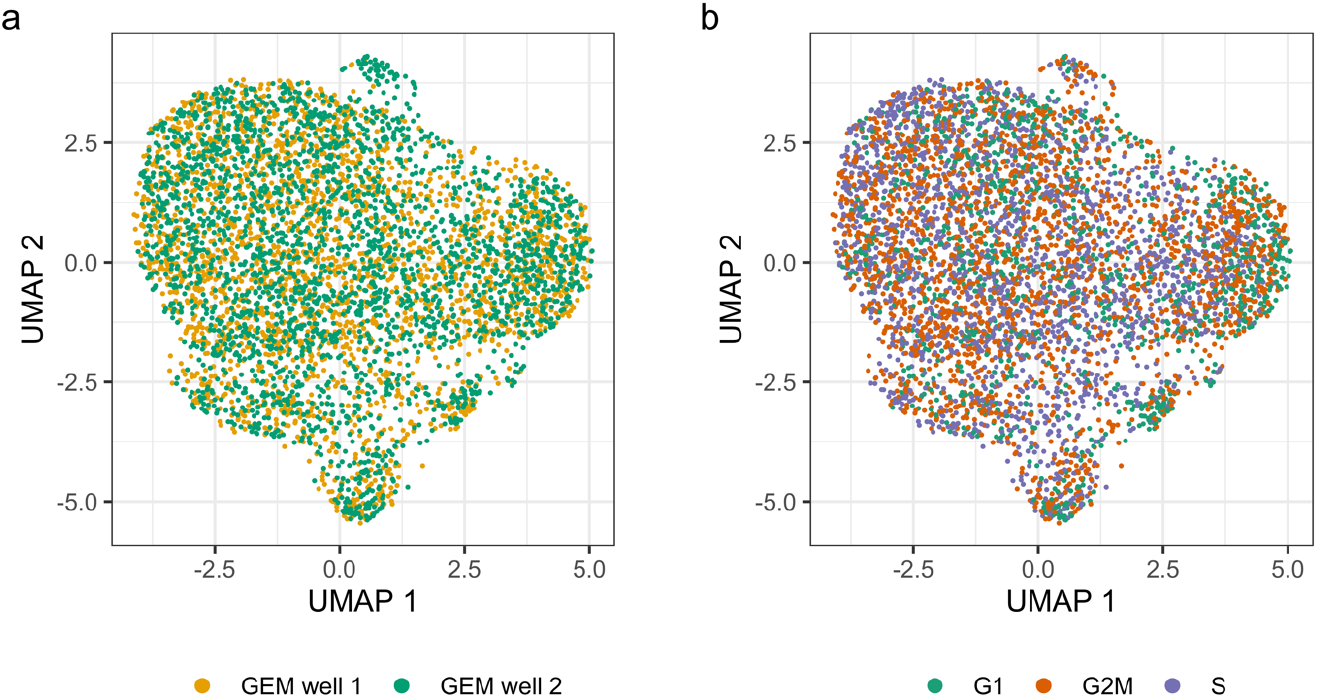
UMAP analysis of single CHO cell gene expression data is not impacted by technical variation from different GEM wells or cell cycle phase. Overlaying the **(a)** source of the scRNA-seq library on the Figure 2b UMAP plot confirms that technical variation introduced across the 2 GEM wells is minimal. The procedure to regress out the effect of cell cycle from the data was successful and **(b)** no clustering according to cell cycle phase was observed.

**Figure S10:**
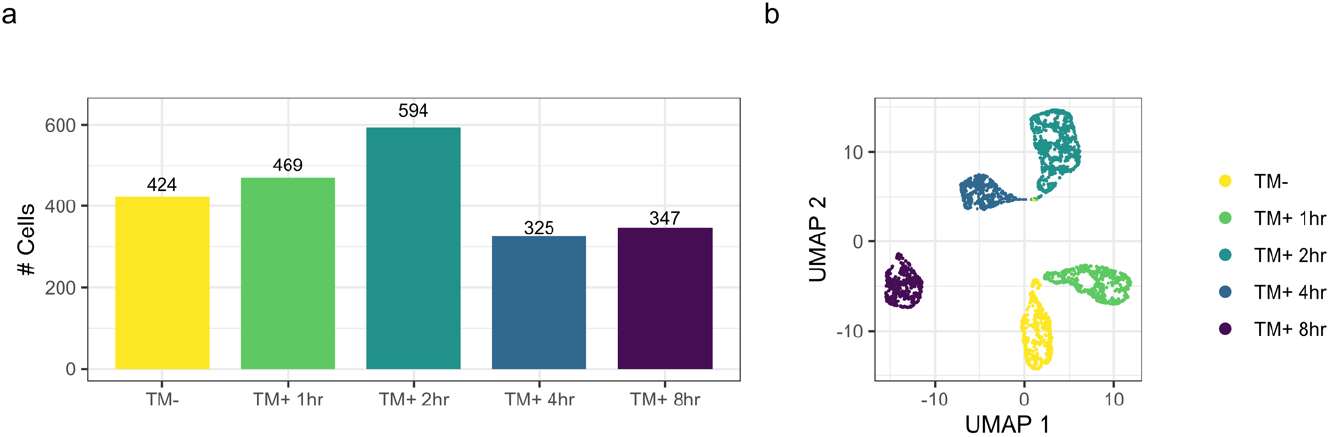
Construction of scWGCNA metacells. To overcome the inherent sparsity of scRNA-seq data for WGCNA the Morabito *et al*. method averages the individual gene expression profiles. For this procedure the SCTransform normalised data aggregated 60 nearest neighbour cells within each condition separately to yield **(a)** 2,159 pseudobulk “metacells”. Using these data, PCA was conducted, and the first 15 principal components (scWGCNA default) were used to generate a **(b)** UMAP plot for the metacells. Each of the 5 samples separate clearly in the UMAP space.

**Figure S11:**
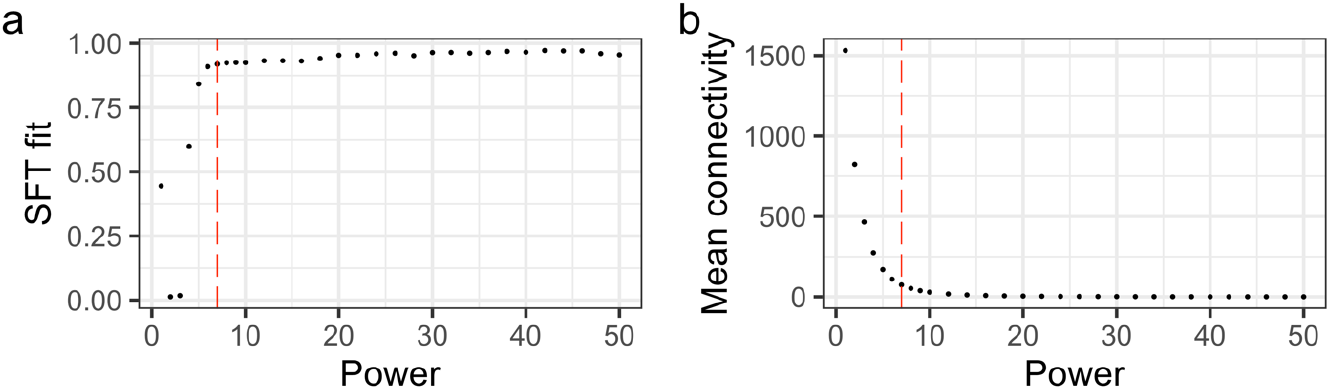
Selection of the WGCNA soft power. A range of power values from 1 to 50 were assessed in terms of the resulting **(a)** scale topology fit (R^2^) and Mean connectivity (k). A power value of 7 was selected to construct the coexpression network (lowest β with a R^2^ > 0.8 and mean k < 100).

**Figure S12:**
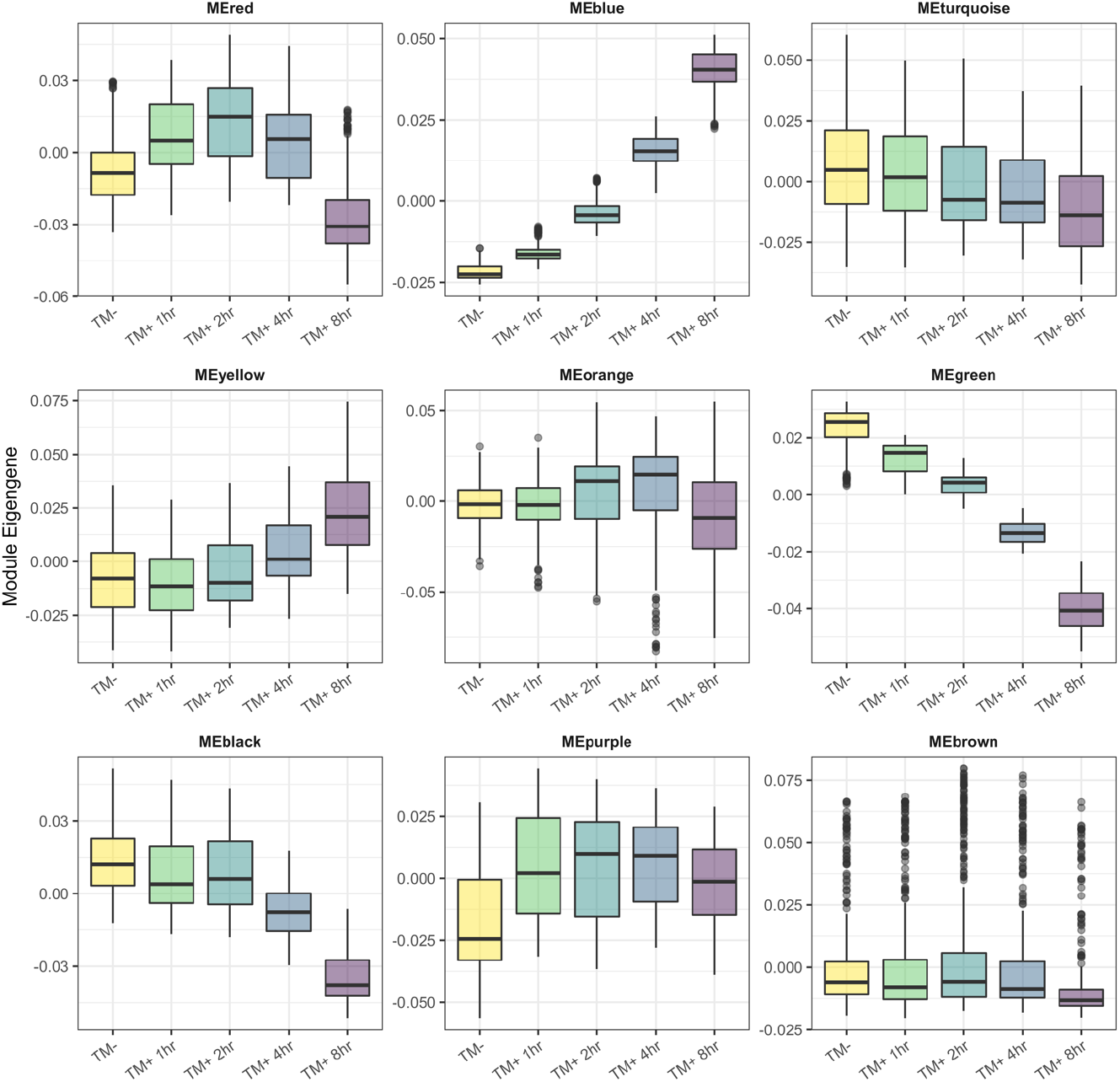
Association of coexpression modules with tunicamycin treatment. PCA was carried out on the metacell dataset for the coexpression modules identified by WGCNA. The values for first principal component, termed the module eigengene (ME), were stratified based on sample identify to assess potential relationships between module eigengene expression and tunicamycin treatment.

**Figure S13:**
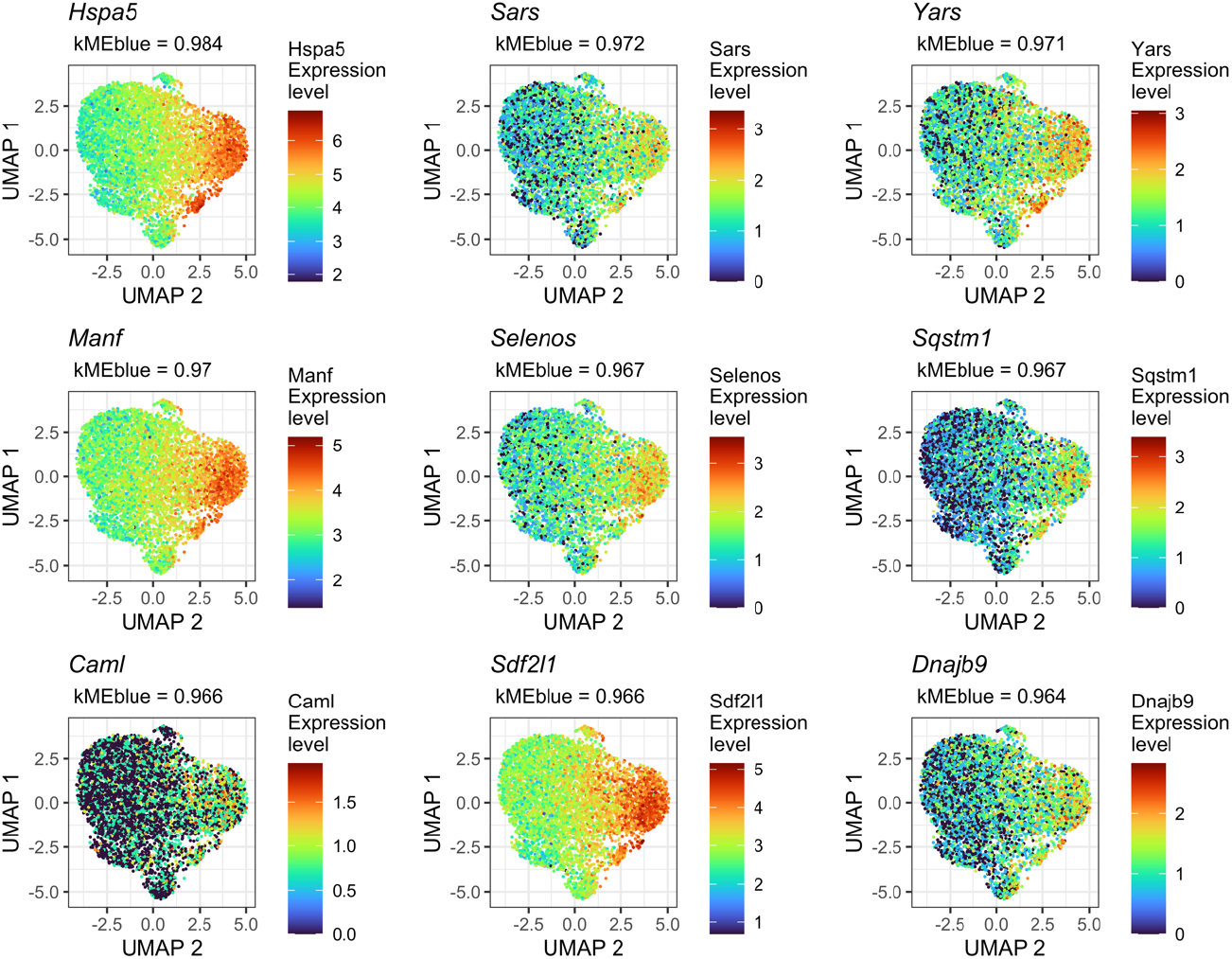
Visualisation of blue coexpression module hub genes. UMAP plots with normalised expression values overlaid illustrate the variability of genes with high intramodular connectivity in the blue module.

**Figure S14:**
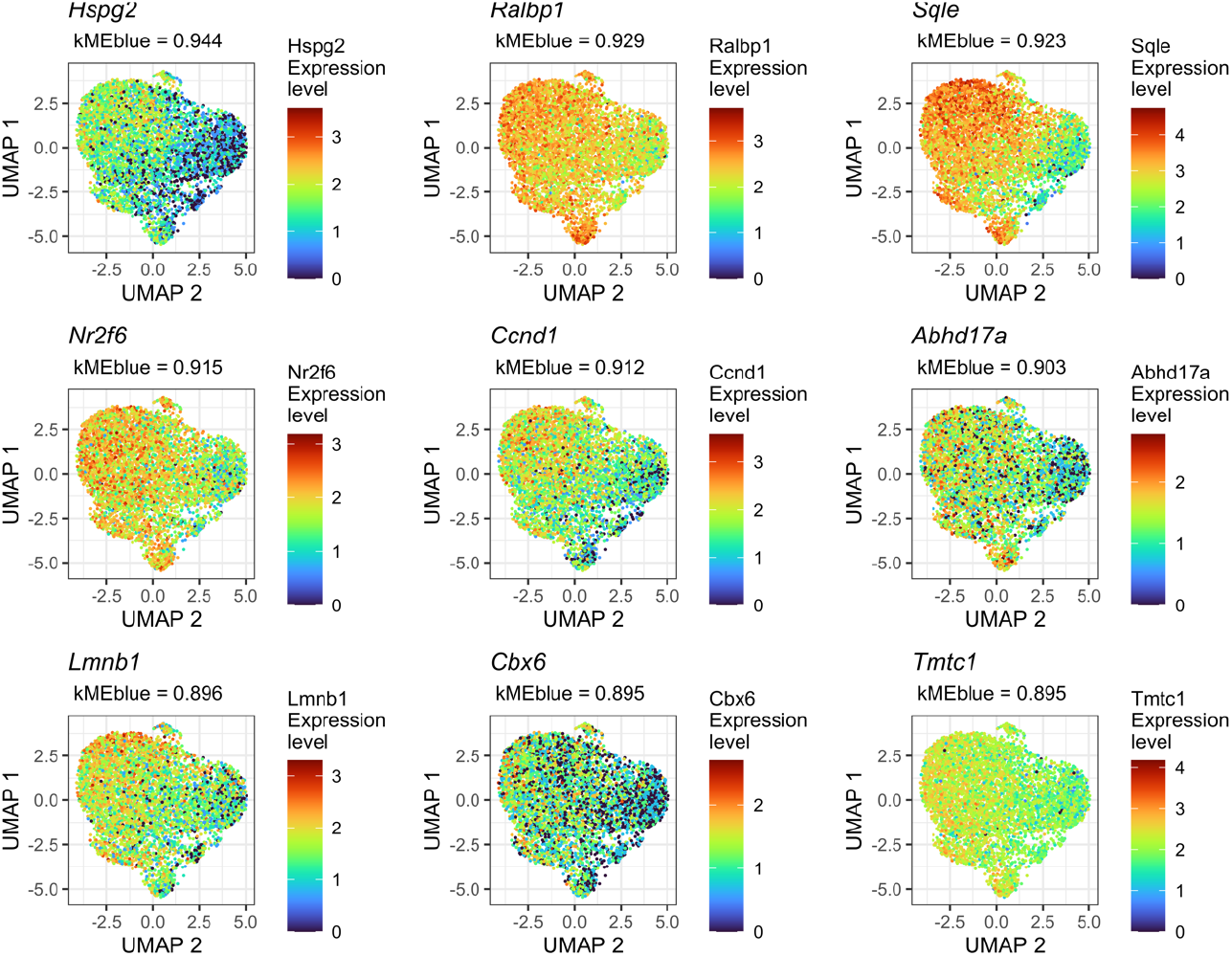
Visualisation of green coexpression module hub genes. UMAP plots with normalised expression values overlaid illustrate the variability of genes with high intramodular connectivity in the green module.

**Figure S15:**
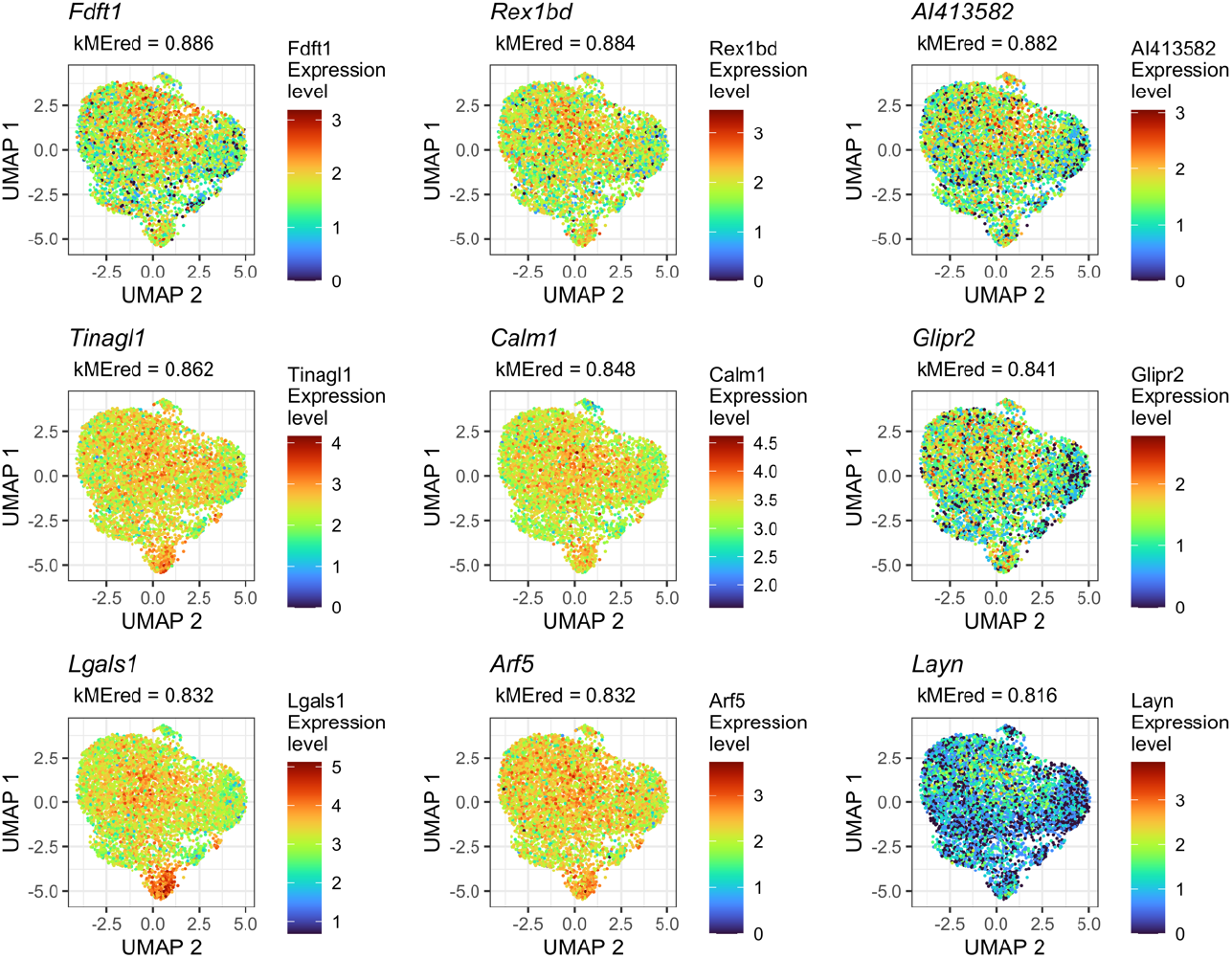
Visualisation of red coexpression module hub genes. UMAP plots with normalised expression values overlaid illustrate the variability of genes with high intramodular connectivity in the red module.

